# Capturing Evolutionary Signatures in Transcriptomes with myTAI

**DOI:** 10.1101/051565

**Authors:** Hajk-Georg Drost, Alexander Gabel, Tomislav Domazet-Lošo, Marcel Quint, Ivo Grosse

**Affiliations:** Sainsbury Laboratory Cambridge, University of Cambridge, Bateman Street, Cambridge CB2 1LR, UK; Martin Luther University Halle-Wittenberg, Institute of Computer Science, Halle (Saale), Germany; Laboratory of Evolutionary Genetics, Division of Molecular Biology, Ruder Boškovic Institute, Zagreb, Croatia; Catholic University of Croatia, Zagreb, Croatia; Department of Molecular Signal Processing, Leibniz Institute of Plant Biochemistry, Halle (Saale), Germany; Martin Luther University Halle-Wittenberg, Institute of Agricultural and Nutritional Sciences, Halle (Saale), Germany; German Center for Integrative Biodiversity Research (iDiv) Halle-Jena-Leipzig, Leipzig, Germany

**Keywords:** Phylostratigraphy, Divergence Stratigraphy, Phylotranscriptomics, Evolution, Development, Evo-Devo

## Abstract

Combining transcriptome data of biological processes or response to stimuli with evolutionary information such as the phylogenetic conservation of genes or their sequence divergence rates enables the investigation of evolutionary constraints on these processes or responses. Such *phylotranscriptomic* analyses recently unraveled that mid-developmental transcriptomes of fly, fish, and cress were dominated by evolutionarily conserved genes and genes under negative selection and thus recapitulated the developmental hourglass on the transcriptomic level. Here, we present a protocol for performing phylotranscriptomic analyses on any biological process of interest. When applying this protocol, users are capable of detecting different evolutionary constraints acting on different stages of the biological process of interest in any species. For each step of the protocol, modular and easy-to-use open-source software tools are provided, which enable a broad range of scientists to apply phylotranscriptomic analyses to a wide spectrum of biological questions.

## Introduction

Transcriptomes carry evolutionary information because expressed genes have different evolutionary ages or are exposed to different selective pressures. In major biological processes such as embryogenesis, metamorphosis, fertilization, senescence, etc. or after biological treatments the set of genes expressed at different stages within these processes varies. Analogously, both environmental and endogenous stimuli elicit responses of different sets of genes. In addition to having varying biological functions, these gene sets may vary regarding their evolutionary signatures like phylogenetic conservation of genes (= gene ages) or their sequence divergence rates (= sequence divergence).

To capture and quantify such evolutionary signatures across development, biological processes or response to stimuli, we developed and established *phylotranscriptomic analyses*, which combine information about gene age and gene sequence divergence with transcriptome data of biological processes and response to stimuli. These analyses allowed the molecular confirmation of the developmental hourglass (Domazet-Lošo and Tautz 2010a), one of the historic principles in evolution and developmental biology originally discovered by von Baer (von Baer 1828).

Moreover, phylotranscriptomic analyses unraveled that the hourglass pattern, which was thought to be a hallmark of animal embryogenesis, is restricted neither to animals nor to embryogenesis. Specifically, such analyses identified molecular hourglass patterns in plants (Quint et al. 2012) and fungi (Cheng et al. 2015) as well as in post-embryonic plant development (Drost et al. 2016).

Phylotranscriptomic analyses are not limited to embryogenesis or other post-embryonic developmental processes. Potential applications of these analyses in other disciplines are studies of life cycles of many different animals, plants, fungi, or bacteria, of metabolic or circadian rhythms, of the mitotic and meiotic cell cycle, of tumor progression, or of a plethora of other fundamental biological processes of life. For example, phylotranscriptomic analyses turned out to be helpful for comparing transcriptomes of three stem cell types in Hydra (Hemmrich et al. 2012) or for reconstructing the evolutionary origin of the neural crest, a *bona fide* innovation of vertebrates (Šestak et al. 2012). In addition, these analyses can be applied to capture evolutionary signatures of temporal responses to different endogenous and exogenous stimuli. Furthermore, phylotranscriptomic analyses can be applied to capture evolutionary signatures on the spatial level in different tissues, organs, cell types, or tumors and allow to study these spatial signatures in development, in other temporal processes, or in response to endogenous and exogenous stimuli.

Despite this great potential, no standardized and reproducible protocol for performing phylotranscriptomic analyses exists to date. To overcome this limitation and to allow a broad range of scientists to capture and quantify evolutionary signatures in transcriptomes by applying such analyses, we developed the open source software packages *<u>createPSmap.pl</u>, orthologr*, and *myTAI* and here provide a user-friendly protocol to apply these tools to any organism and biological study of interest.

## Methodological Background

Given the wide variety of possible applications of phylotranscriptomic analyses, we focus on development as an example use case of our protocol.

Evolutionary signatures of transcriptomes can be captured by computing transcriptome indices at different measured stages of development, combining these computed values to a transcriptome index profile across the measured stages, and comparing this profile with a flat line. A profile not significantly deviating from a flat line indicates the absence of significant variations of the computed transcriptome index from stage to stage. In contrast, a profile significantly deviating from a flat line indicates the presence of significant variations from stage to stage. We refer to any transcriptome index profile significantly deviating from a flat line as phylotranscriptomic pattern or evolutionary signature.

The computation of the *transcriptome age index (TAI*) (Domazet-Lošo and Tautz 2010a) (Supplementary Information) requires the determination of the evolutionary ages of the genes of the studied species. For this purpose, we developed a procedure termed *phylostratigraphy* (Domazet-Lošo et al. 2007), which briefly described works as follows: First, a sequence homology search of the proteins of the studied species against a database of proteins of completely sequenced genomes from species covering all kingdoms of life is performed. Second, these species are sorted into sets named *phylostrata* (PS) corresponding to hierarchically ordered phylogenetic nodes along the tree of life. PS 1 denotes the set of all living species, PS 2 denotes the set of species of the same domain as the query species, PS 3 denotes the set of species of the same kingdom as the query species, etc., and the highest PS denotes the set consisting of only the studied species. Third, each protein of the studied species is assigned to the lowest PS in which at least one homolog with a predefined threshold on the degree of homology was detectable. The resulting assignment of one PS to each protein of the studied genome is called *phylostratigraphic map*, and the PS of a given protein or protein-coding gene is often loosely called *gene age*.

The TAI at a given stage of development is then obtained by joining this phylostratigraphic map with expression data at that stage and by computing the weighted mean of the PS, where the weights are the stage-specific expression levels (Supplementary Information) (Domazet-Lošo and Tautz 2010a). Loosely speaking, the TAI at a given stage of development is the mean evolutionary age of the genes expressed at that stage, and the TAI profile is the profile of these mean ages across different stages of development. Stages with high TAI values are stages where evolutionarily old genes (in low PS) are more lowly expressed - and evolutionarily young genes (in high PS) are more highly expressed - than in other stages.

The computation of the *transcriptome divergence index (TDI*) (Quint et al. 2012, Drost et al. 2015) (Supplementary Information) requires the determination of the degree of selection of the genes of the studied species. For this purpose, we developed a procedure termed *divergence stratigraphy* (Quint et al. 2012, Drost et al. 2015), which works analogously to phylostratigraphy as follows: First, a set of orthologs of the proteins of the studied species is obtained in a closely related species. Second, rates of non-synonymous (dN) and synonymous (dS) substitutions as well as the dN/dS ratio are estimated for each pairwise alignment of the protein-coding genes of the protein of the studied species and its ortholog. Third, the continuous dN/dS ratios are discretized by sorting them into e. g. 10 groups of equal sizes, where group 1 contains the genes with the lowest dN/dS ratios, and group 10 contains the genes with the highest dN/dS ratios. In analogy to PS, these groups of genes of similar dN/dS ratios are called *divergence strata (DS)*. The resulting assignment of one DS to each protein of the studied species with an ortholog is called *divergence stratigraphic map* (Quint et al. 2012, Drost et al. 2015), and the DS of a given protein-coding gene is often loosely called *sequence divergence* denoting the evolutionary rate of a protein (Zhang and Yang 2015).

The TDI at a given stage of development is then obtained in analogy to obtaining the TAI by joining this divergence stratigraphic map with expression data at that stage and by computing the weighted mean of the DS (Quint et al. 2012, Drost et al. 2015). Loosely speaking, the TDI at a given stage of development is the mean sequence divergence of the genes expressed at that stage, and the TDI profile is the profile of these mean sequence divergences across different stages of development. Stages with high TDI values are stages where genes under strong negative selection (in low DS) are more lowly expressed - and genes under weaker negative selection or even positive selection (in high PS) are more highly expressed - than in other stages.

To assess the significance of deviations of transcriptome index profiles from a flat line, we proposed the *flat-line test* (Quint et al. 2012; Drost et al. 2015). The *flat-line test* is a permutation test that randomly assigns PS or DS to the genes of investigation. This random assignment is used to compute TAI or TDI profiles and produces random patterns of transcriptome conservation. Subsequently, the variance is then used as a measure to quantify the variation of these random transcriptome indices between stages. This procedure is performed independently 10,000 times and the resulting variance values are compared with the actual variance of TAI or TDI patterns. If the value of the actual TAI or TDI variance value is less than 0.05 we declare the corresponding pattern as significantly deviating from a flat line and as not deviating from a flat line otherwise (Supplementary Information).

Two phylotranscriptomic patterns of particular interest in plant and animal embryogenesis are the hourglass pattern and the early-conservation pattern. To test the presence or absence of these patterns we introduced the *reductive hourglass test* and *reductive early-conservation test* (Drost et al. 2015). The reductive hourglass test quantifies the degree of agreement of the transcriptome index profile to an hourglass-like high-low-high pattern, while the reductive early-conservation test quantifies the degree of agreement with an early conservation-like low-low-high pattern. In addition, we designed *myTAI* such that users can easily build customized statistical tests for assessing the significance of any pattern deviating from a flat line (Supplementary Information).

To further scrutinize observed phylotranscriptomic patterns we introduced relative expression profiles per PS and per DS (Domazet-Lošo and Tautz 2010a; Drost et al. 2015) (Supplementary Information). Relative expression levels allow the visualization of the average expression behavior of genes from the same PS or the same DS across the biological process of interest. Specifically, the mean expression profile of each PS and each DS across all stages is linearly transformed to a normalized profile ranging from 0 to 1, and the resulting normalized profile is called *relative expression profile* of the corresponding PS and DS.

## Protocol Applications

As introduced before, phylotranscriptomic analyses enabled us to unravel the existence of phylotranscriptomic hourglass patterns in animals (Domazet-Lošo and Tautz 2010a), plants (Quint et al. 2012; Drost et al. 2015; Drost et al. 2016), and fungi (Cheng et al. 2015). These findings suggested that ontogenetic processes occurring during development are related to phylogenetic processes occurring during evolution (Von Baer 1828; Sander 1983; Duboule 1994; Richardson 1995; Raff 1996; Richardson et al. 1997; Richardson 1999; Hazkani-Covo et al. 2005; Irie and Sehara-Fujisawa 2007; Artieri et al. 2009; Cruickshank and Wade 2008; Kalinka et al. 2010; Domazet-Lošo and Tautz 2010a; Yanai et al. 2011; Irie and Kuratani 2011; Levin et al. 2012; Willmore 2012; Svorcová 2012; Tian et al. 2013; Wang et al. 2013; Gerstein et al. 2014; Levin et al. 2016; Gossmann et al. 2016).

Specifically, it has been found that the pattern of morphologically dissimilar-similar-dissimilar embryos between related animal species, the so-called developmental hourglass pattern (Duboule 1994; Raff 1996), is mirrored by a similar hourglass-like high-low-high pattern of TAI and TDI profiles on the phylotranscriptomic level (Domazet-Lošo and Tautz 2010a; Drost et al. 2015). Moreover, it has been found that the stage or period of maximum transcriptome conservation during mid embryogenesis coincides with the morphological stage of maximum conservation defined as phylotypic stage (Sander 1983) or phylotypic period (Richardson 1995; Raff 1996; Richardson et al. 1997). In this context, phylotranscriptomic analyses have provided a molecular explanation for and thus deepened our understanding of the relation between evolution and development, and we believe that such analyses will advance current and future evo-devo research, too.

However, the applicability of the protocol and the developed software packages is not restricted to the study of development. As specified in section *Methodological Background*, this protocol can be applied to phylotranscriptomic analyses of any transcriptome data set of any biological study of any species. In addition to phylotranscriptomic analyses, individual modules of this protocol can be used for performing different analyses independently of each other.

For example, phylostratigraphy alone has previously been performed to detect orphan genes (Tautz and Domazet-Lošo 2011) or the evolutionary origin of specific classes of genes such as cancer genes (Domazet-Lošo and Tautz 2010b) or transcription factors (De Mendoza et al. 2013). Likewise, divergence stratigraphy can simply be performed for quantifying the rate of synonymous versus non-synonymous substitution rates of protein-coding genes in a genome of choice (Drost et al. 2015).

The modularity of the protocol also allows users to use their own modules for phylostratigraphy or divergence stratigraphy. This modularity for example, enables to study the influence of different phylostratigraphies or divergence stratigraphies on TAI or TDI profiles. In particular, the influence of potential underestimations of gene ages by BLAST approaches (Moyers and Zhang 2015, 2016) can be systematically investigated using this protocol.

In general, this protocol was designed to make it easily applicable for life scientists. For this purpose, we provide software tools and step-by-step instructions for every part of this protocol. For example, we provide the Perl script *<u>createPSmap.pl</u>* for performing BLAST searches and computing phylostratigraphic maps, the R package *orthologr* for identifying orthologs and computing divergence stratigraphic maps, and the R package *myTAI* for computing TAI and TDI profiles across developmental stages, for performing statistical tests such as the flat-line test, for computing relative expression profiles for all PS and DS, or for producing scientific visualizations of observed phylotranscriptomic patterns in publication quality. All parts of the protocol are demonstrated by using the same example data set covering seven stages of *A.thaliana* embryo development (Quint et al. 2012; Drost et al. 2015).

All software tools are publicly available under an open source license (https://github.com/AlexGa/Phylostratigraphy, https://github.com/HajkD/orthologr, and https://github.com/HajkD/myTAI). An extensive documentation of each function as well as six tutorials covering different phylotranscriptomic analyses are part of the *orthologr* package, the *myTAI* package, and the Supplementary Information.

Furthermore, pre-computed phylostratigraphic maps and divergence stratigraphic maps can be obtained from a public repository (https://github.com/HajkD/published_phylomaps). In this way, users can easily adapt the provided protocol to the organism, biological process, and data sets of their interest with minimal computational effort.

## Comparison with other similar techniques

Possible alternatives to phylostratigraphy are based on Wagner-Parsimony or Phylogenetic-Reconciliation (Capra et al. 2013) and are summarized in the software tool ProteinHistorian (Capra et al. 2012; Capra et al. 2013). ProteinHistorian allows performing alternative gene age assignment methods that can then be used by *myTAI* to compute TAI profiles and to perform all other analyses based on phylostratigraphic maps resulting from these alternative methods for estimating gene ages.

Possible alternatives to divergence stratigraphy are based on phylogenetic inference. Here, the metaPhOrs repository (http://orthology.phylomedb.org/) can be accessed to retrieve pre-computed phylogeny-based orthology predictions that can then be used to estimate sequence substitution rates. Alternative methods for estimating substitution rates are provided by the R package *orthologr*, where the function dNdS() allows users to choose a variety of alternative methods for estimating substitution rates of predicted orthologs (for detailed information see https://github.com/HajkD/orthologr/blob/master/vignettes/dNdS_estimation.Rmd). These alternative divergence stratigraphic maps can then be used by *myTAI* for computing TDI profiles and for performing all other analyses.

A broadly applied alternative approach for associating developmental transcriptomes with evolutionary constraints is comparative transcriptomics, which has been used to study developmental hourglass patterns in several species (Kalinka et al. 2010; Irie and Kuratani 2011; Levin et al. 2012; Romero et al. 2012; Wang et al. 2013; Dunn et al. 2013; Warnefors and Kaessmann 2013; Gerstein et al. 2014; Nesculea and Kaessmann 2014; Levin et al. 2016). Comparative transcriptomics analyzes gene expression diversity between orthologs of two or more species based on the empirical finding that gene expression diversity of orthologs correlates with developmental dissimilarity (Roux et al. 2015).

## Limitations of the Protocol

One limitation of the protocol is that it can only be applied to species for which protein sequences are annotated and for which this annotation matches the transcriptome annotation of the corresponding gene expression data set.

A second limitation is that computing a phylostratigraphic map can take several hours, several days, or even several weeks depending on the number of query sequences and the size of the database of proteins of completely sequenced genomes and may thus require a computing cluster.

A third limitation is that users interested in applying a custom taxonomy to ***createPSmap.pl.*** need to perform additional steps to retrieve a customized phylostratigraphic map.

However, as the ***myTAI*** package is designed to take any custom phylostratigraphic map or divergence stratigraphic map as input, all of the subsequent phylotranscriptomic analyses of this protocol can be performed irrespectively of the origins of the phylostratigraphic or divergence stratigraphic maps (Supplementary Information).

## Example Experiment

The following example protocol is divided into four conceptual parts. The first part (steps 1-2) covers the construction of phylostratigraphic and divergence stratigraphic maps. The second part (steps 3-10) includes the computation and visualization of TAI and TDI profiles. The third part (steps 11-12) covers the application of three statistical tests for quantifying the significance of the observed phylotranscriptomic patterns, and the fourth part (steps 13-18) includes the computation and visualization of relative expression profiles (Fig. 1).

**Figure 1.**
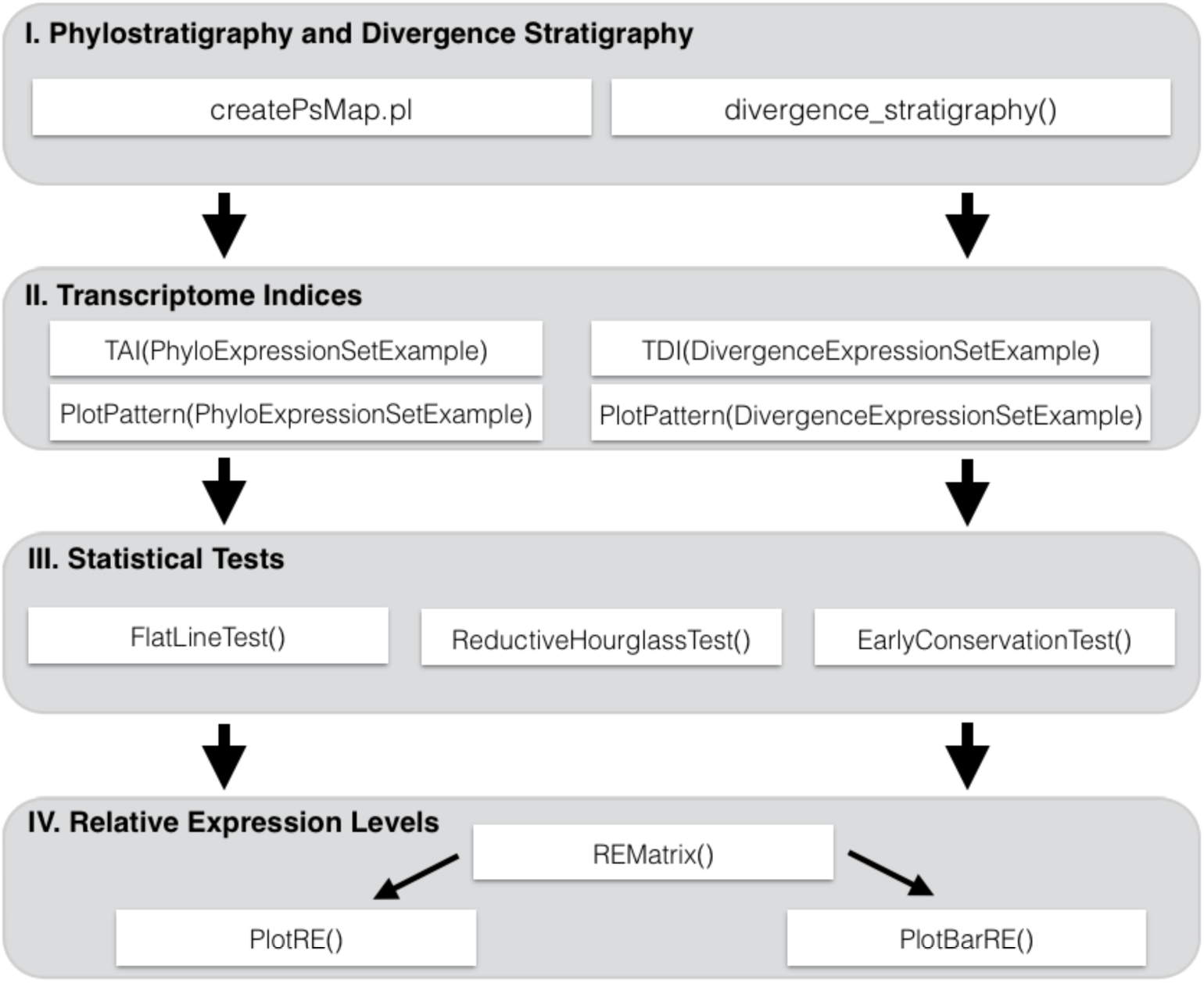
Flow Chart. A step-by-step instruction illustrating the workflow of the protocol.

In step 1 of the protocol, the Perl script *createPSmap.pl.* is used for computing the phylostratigraphic map (Fig. 2). The input files to *createPSmap.pl.* are *a fasta* file of protein sequences of the studied species of and a database of proteins of completely sequenced genomes in *fasta* format. Phylogenetic information is provided in the header of *fasta* sequences and can be customized by following the tutorial at https://github.com/AlexGa/Phylostratigraphy. The output of *createPSmap.pl.* is the phylostratigraphic map, i.e., a table storing the gene id and the PS of each protein-coding gene of the studied species (https://github.com/AlexGa/Phylostratigraphy).

**Figure 2.**
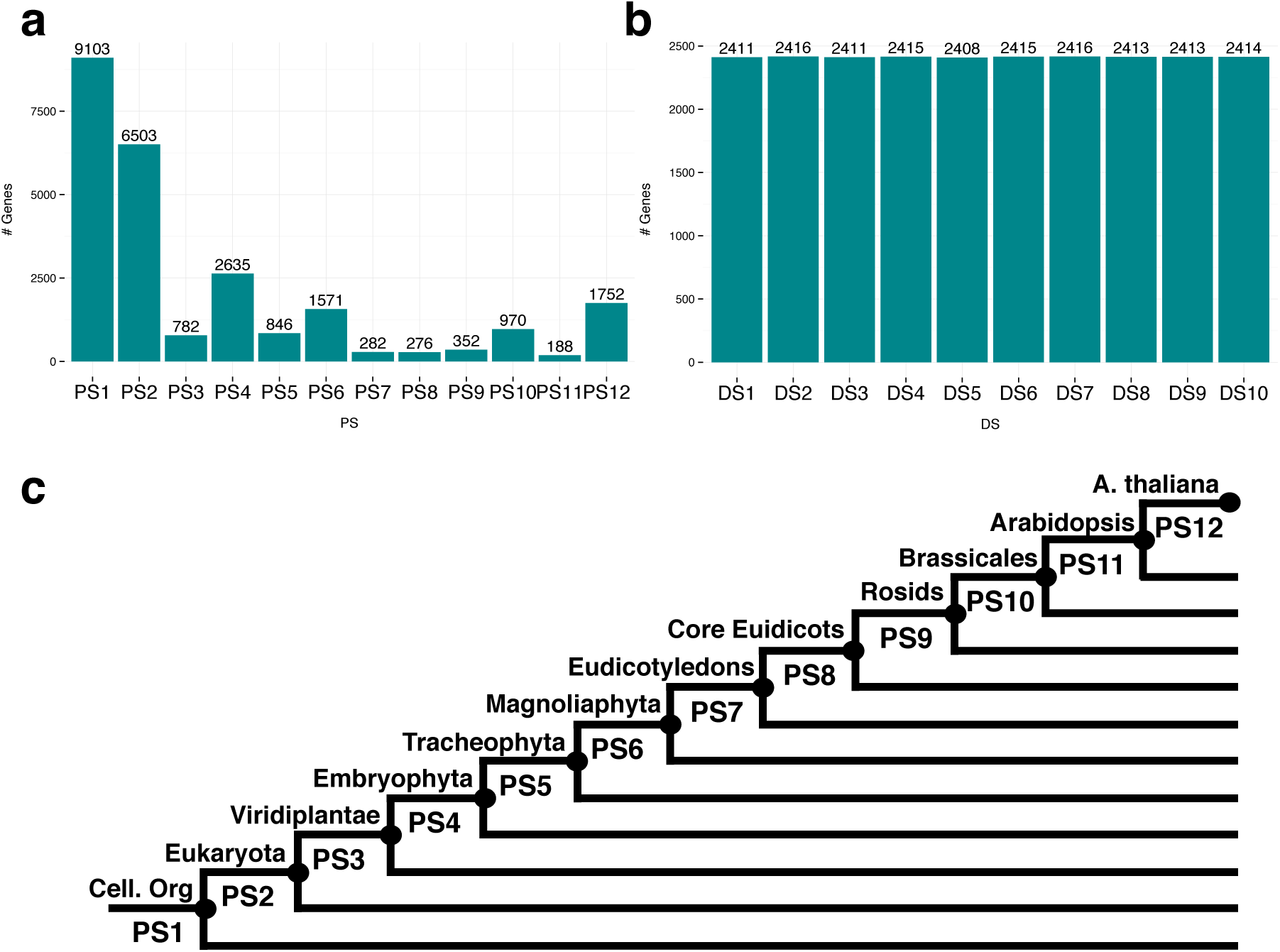
Histograms of phylostrata and divergence strata and phylostratigraphic map. PS histogram (a) and DS histogram (b) of *A.thaliana* genes. The uniform DS histogram is due to the definition as deciles, **c** Phylostrati graphic map of *A.thaliana*. The most distant taxonomic categoiy is PS1 [cellular organisms) and the closest categoiy is PS12 (*A.thaliana*).

In step 2, function divergence_stratigraphy() of R package *orthologr* is used for computing the divergence stratigraphic map. In the example data set, the arguments *query_file* and *subject_file* refer to the downloaded CDS files *Athaliana_167_cds.fa* and *Alyrata_107_cds.fa* (see *Prerequisite tools* for details) and denote the input files of function divergence_stratigraphy(). The output of the function divergence_stratigraphy() is the divergence stratigraphic map, i.e. a table storing the gene id and the DS of each protein-coding gene of the studied species that has an ortholog in the other species (https://github.com/HajkD/orthologr/blob/master/vignettes/divergence_stratigraphy.Rmd).

In step 3, the phylostratigraphic map and the divergence stratigraphic map are matched with a transcriptome data set covering the studied biological process. This step is accomplished by function *MatchMap()* of R package *myTAI*, which takes a phylostratigraphic map or divergence stratigraphic map and an expression data set as input and returns a table storing the gene id, its PS or DS, and its expression profile as output. We denote this output data as *PhyloExpressionSet* or *DivergenceExpressionSet*.

Phylostratigraphy and divergence stratigraphy need to be performed only once for each species, and phylostratigraphic maps and divergence stratigraphic maps are available for a variety of species at https://github.com/HajkD/published_phylomaps.

In step 4, R package *myTAI* and pre-formatted data sets required for subsequent analyses of the protocol are loaded into the current R session.

Steps 5 and 6 visualize the phylostratigraphic map (Fig. 2a) and the divergence stratigraphic map (Fig. 2b) by plotting histograms of absolute or relative frequencies of genes per PS or per DS, respectively.

TAI and TDI profiles are computed and visualized in steps 7-10 (Fig. 3). Both functions TAI() and TDI() take a *PhyloExpressionSet* or *DivergenceExpressionSet* as input and compute the TAI and TDI values as output. These profiles are then visualized by function PlotPattern().

**Figure 3.**
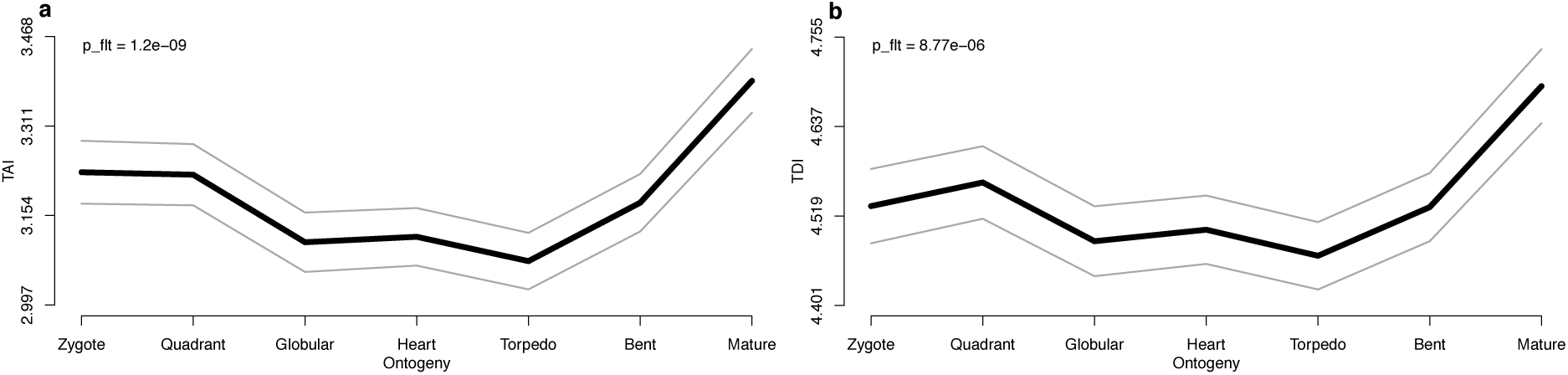
Transcriptome indices of *A. thaliana* embryogenesis. Visualization of (**a**) the TAI profile and (**b**) the TDI profile covering seven stages of *A.thaliana* embryogenesis. Grey lines represent the standard deviation of randomly permuted TAI or TDI profiles. The statistical significance of observed patterns (*p*-values) are computed using the *flat-line test* (Drost et al. 2015).

In steps 11 and 12, *p*-values of the TAI and TDI profiles are computed according to the *flat-line test*, the *reductive hourglass test* (11.A), or the *reductive early-conservation test* (11.B). The flat-line test assesses the deviation of a TAI or TDI profile from a flat line, while the reductive hourglass test and the reductive early-conservation test indicate the presence of a high-low-high pattern or a low-low-high pattern, respectively, based on an *a-priori* definition of early, mid (phylotypic), and late phases of development (Drost et al. 2015) (Supplementary Information).

In steps 13-16, tables of relative expression profiles for each PS and each DS are obtained and visualized by using functions REMatrix() and PlotRE() of R package *myTAI*.

In steps 17-18, a group-specific visualization of relative expression levels is performed by function PlotBarRE() of R package *myTAI*, and potential differences between groups are assessed by a Kruskal-Wallis rank sum test (Fig. 4).

**Figure 4.**
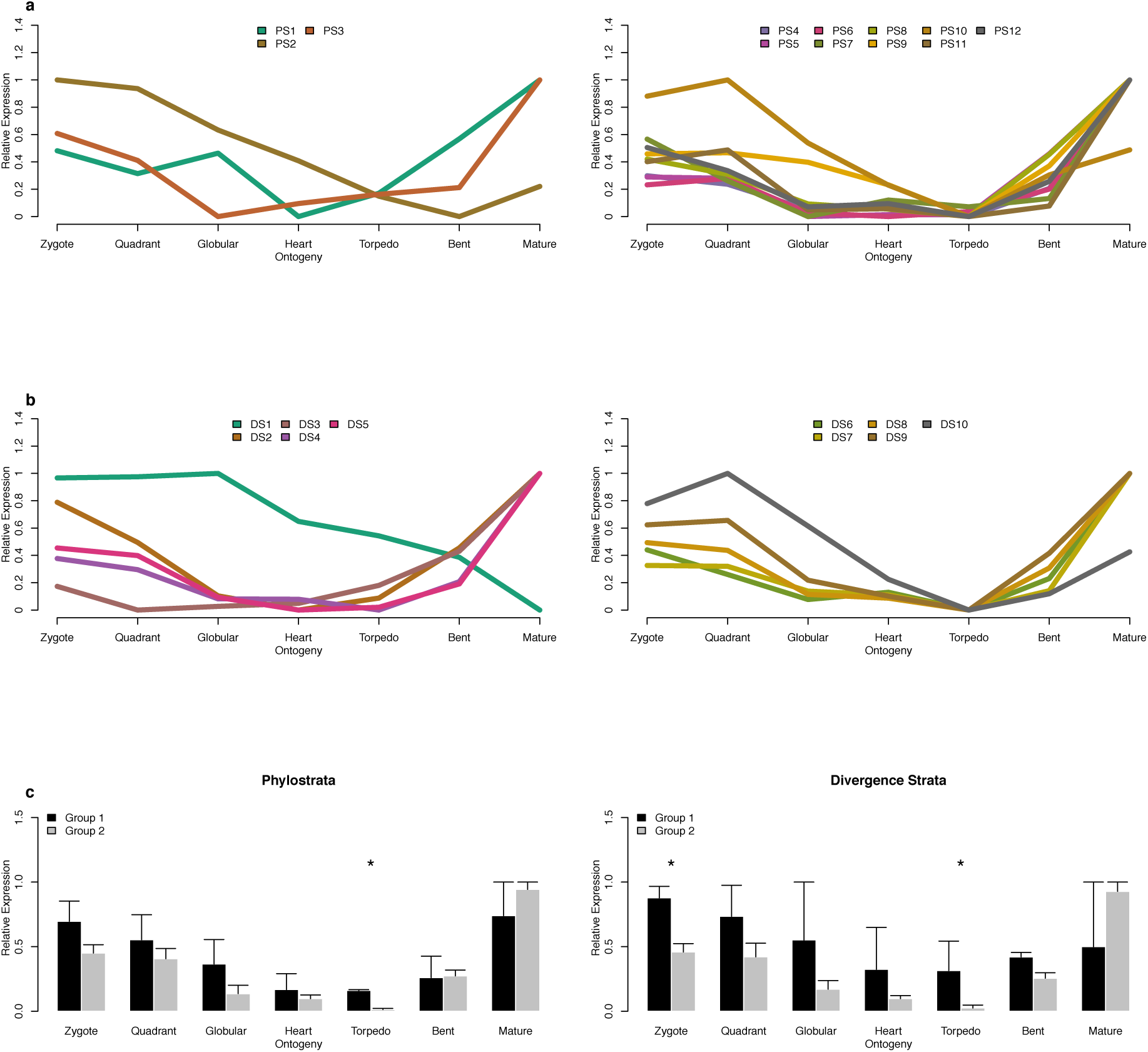
Relative expression profiles of *A.thaliana* embryogenesis. Relative expression profiles of twelve PS (**a**) covering seven stages of *A.thaliana* embryogenesis. PS are divided into two groups to analyze co-expression patterns of PS before (PS 1-3) and after (PS 4-12) the emergence of embryogenesis in plant evolution. Relative expression profiles often DS (**b**) of the same stages. DS are divided into two groups (DS 1-2 versus DS 3-10) to analyze co-expression patterns of DS with highly negative (purifying) selection (DS 1-2) and more relaxed or even positive selection (DS 3-10). **c**. Bar plots of mean relative expression levels for PS and DS groups. *p*-values of the difference of mean relative expression levels between PS groups or DS groups are obtained by a Kruskal-Wallis rank-sum test. Developmental stages with significant differences of mean relative expression levels are marked by asterisks (Drost et al. 2015).

## Prerequisite tools

### BLAST

The Basic Local Alignment Search Tool (BLAST) (Altschul et al. 1990) is used to determine gene homology relationships in phylostratigraphy and divergence stratigraphy and can be downloaded from ftp://ftp.ncbi.nlm.nih.gov/blast/executables/blast+/LATEST/. A detailed installation guide can also be found at https://github.com/HajkD/orthologr/blob/master/vignettes/Install.Rmd#install-blast.

### Perl and Java Programming Environments

The Perl programming language can be downloaded from https://www.perl.org/ and the Java compiler and interpreter can be obtained from https://www.java.com/de/download/.

The pipeline to perform phylostratigraphy can be downloaded from https://github.com/AlexGa/Phylostratigraphy.

### R Programming Environment

The R programming environment (R Core Team 2016) can be downloaded from http://cran.r-project.org/ and installed under Linux, Mac OSX, or Windows.

The R package *orthologr* can be downloaded and installed from https://github.com/HajkD/orthologr.

The R package *myTAI* can be downloaded and installed from https://github.com/HajkD/myTAI.

### R Package Dependency

*myTAI* (Drost 2016) uses the following open source packages from CRAN: Rcpp (Eddelbuettel and Francois 2011), nortest (Gross and Ligges 2014), fitdistrplus (Delignette-Muller and Dutang 2015), doParallel (Weston 2014), dplyr (Wickham and Francois 2015), RColorBrewer (Neuwirth 2014), taxis (Chamberlain and Szocs 2013), ggplot2 (Wickham 2009), and edgeR (Robinson et al. 2010).

*orthologr* (Drost et al. 2015) uses the following open-source packages from CRAN and Bioconductor (Huber et al. 2015): Rcpp (Eddelbuettel and Francois 2011), doParallel (Weston 2014), dplyr (Wickham and Francois 2015), seqinr (Charif and Lobry 2007), data.table (Dowle 2014), Biostrings (Pages et al. 2007), RSQLite (Wickham et al. 2014), stringr (Wichham 2015), IRanges (Lawrence et al. 2013), DBI (R Special Interest Group on Databases 2014), and S4Vectors (Pages et al. 2014).

### orthologr and myTAI

After installing R packages *orthologr* and *myTAI*, they can be loaded into the current R session by commands

*library(orthologr)*

*library(myTAI)*

### Data

To perform phylostratigraphy, the reference genome database needs to be downloaded from http://msbi.ipb-halle.de/download/phyloBlastDB_Drost_Gabel_Grosse_Quint.fa.tbz. This database currently stores 17,582,624 amino acid sequences covering 4,557 species. After downloading file phyloBlastDB, it can be unpacked by opening a terminal application and typing

> *tar xfvj phyloBlastDB_Drost_Gabel_Grosse_Quint.fa.tbz*

The header of the FASTA-files of the studied species (e.g. Athaliana_167_protein.fa) must fulfill the following specification

> >GeneID | [species_name] | [taxonomy]

The corresponding taxonomy starts with super kingdoms limited to Eukaryota, Archaea, and Bacteria. For the example data set this yields:

> >ATCG00500.1|PACid:19637947 | [Arabidopsis thaliana] | [Eukaryota; Viridiplantae; Streptophyta; Streptophytina; Embryophyta; Tracheophyta; Euphyllophyta; Spermatophyta; Magnoliophyta; eudicotyledons; core eudicotyledons; rosids; malvids; Brassicales; Brassicaceae; Camelineae; Arabidopsis]

To download the coding sequence (CDS) files of *A.thaliana* and *A.lyrata* from the Phytozome database (Goodstein et al. 2012), the following R command-line tool can be used:

> # download the CDS file of *A. thaliana download.file (url = "ftp://ftp.jgi-psf.org/pub/compgen/phytozome/v9.0/Athaliana/annotation/Athaliana_167_cds.fa.gz", <u>destfile = "Athaliana_167_cds.fa.gz"</u>)*
>
> # download the CDS file of A.lyrata *download.file (url = "ftp://ftp.jgi-psf.org/pub/compgen/phytozome/v9.0/Alyrata/annotation/Alyrata_107_cds.fa.gz", <u>destfile = "Alyrata_107_cds.fa.gz"</u>)*

Next, the files Athaliana_167_cds.fa.gz and Alyrata_107_cds.fa.gz need to be unpacked. These CDS files are then used by R package *orthologr* for performing divergence stratigraphy.

*myTAI* includes two example data sets, each containing a gene-expression time course covering seven stages of *A. thaliana* embryogenesis starting from the zygote stage to mature embryos as well as a phylostratigraphic map and divergence stratigraphic map of each protein-coding gene of *A.thaliana* (Xiang et al. 2011; Quint et al. 2012) with an ortholog in *Arabidopsis lyrata* (Drost et al. 2015). These example data sets are loaded into the current R session by commands

> *data(PhyloExpressionSetExample)*
>
> *data(DivergenceExpressionSetExample)*

The structure of these example data sets can be displayed by commands

> *head(PhyloExpressionSetExample, 3)*
>
> *head(DivergenceExpressionSetExample, 3)*

The structure of the data set resembles a standard format for all functions and procedures implemented in *myTAI*. To distinguish data sets storing phylostratigraphic maps combined with gene expression data and sequence divergence stratigraphic maps combined with gene expression data, the notation *PhyloExpressionSet* and *DivergenceExpressionSet* is used in the documentation *of myTAI*.

### Protocol Steps

1. Compute phylostratigraphic map:

> *perl <u>createPSmap.pl</u>—organism Athaliana_167_protein_with_new_Header.fa*
>
> *— database phyloBlastDB_Drost_Gabel_Grosse_Quint.faphyloBlastDB.fa*
>
> *— evalue 1e-5—threads 64—blastPlus*

2. Compute divergence stratigraphic map with reciprocal best hit:

> *library(orthologr)*

\# compute the divergence stratigraphic map of *A.thaliana* vs. *A.lyrata*

> *Ath_vs_Aly_DM <- divergence_stratigraphy(*
>
> *query_file = "Athaliana_167_cds.fa"*,
>
> *subject_file = "Alyrata_107_cds.fa"*,
>
> *eval = "1E-5"*,
>
> *ortho_detection = "RBH"*,
>
> *comp_cores = 1)*

A. Compute divergence stratigraphic map with best hit:

> *library(orthologr)*

\# compute the divergence map of *A.thaliana* vs. *A.lyrata using BLAST best hit*

> *Ath_vs_Aly_DM <- divergence_stratigraphy(*
>
> *query_file = "Athaliana_167_cds.fa"*,
>
> *subject_file = "Alyrata_107_cds.fa"*,
>
> *eval = "1E-5"*,
>
> *ortho_detection = "BH"*,
>
> *comp_cores = 1)*

3. Match phylostratigraphic map of step 1 or divergence stratigraphic map of step 2 with an expression data set:

> *library(myTAI)*
>
> *PhyloExpressionSet <- MatchMap(PhyloMap, ExpressionMatrixExample)*
>
> *DivergenceExpressionSet <- MatchMap(DivergenceMap, ExpressionMatrixExample)*

4. Load R package *myTAI* and read data of step 3 into the current R session:

> *library(myTAI)*
>
> *data(PhyloExpressionSetExample)*
>
> *data(DivergenceExpressionSetExample)*

A. Load R package *myTAI* and read a custom expression data set from a hard drive:

> *library(myTAI)*
>
> *myPhyloExpressionSet <- read.csv(“PhyloExpressionSet.csv”, sep = “,”, header = TRUE)*
>
> *myDivergenceExpressionSet <- read.csv(“DivergenceExpressionSet.csv”, sep = “,”, header = TRUE)*

5. Visualize the phylostratigraphic map of step 1 (Fig. 2a):

> *PlotDistribution (PhyloExpressionSet = PhyloExpressionSetExample*,
>
> *xlab = “Phylostratum”)*

6. Visualize the divergence stratigraphic map of step 2 (Fig. 2b):

> *PlotDistribution (PhyloExpressionSet = DivergenceExpressionSetExample*,
>
> *xlab = “Divergence stratum”)*

7. Compute the TAI profile from the data set of step 3:

> *TAI(PhyloExpressionSetExample)*

8. Visualize the TAI profile from step 7 (Fig. 3a):

> *PlotPattern (ExpressionSet = PhyloExpressionSetExample*,
>
> *type = “l”*,
>
> *lwd = 6*,
>
> *xlab = “Ontogeny”*,
>
> *ylab = “TAI”)*

9. Compute the TDI profile from the data set of step 3:

> *TDI(DivergenceExpressionSetExample)*

10. Visualize the TDI profile from step 9 (Fig. 3b):

> *PlotPattern (ExpressionSet = DivergenceExpressionSetExample, type = “l”*,
>
> *lwd = 6*,
>
> *xlab = “Ontogeny”*,
>
> *ylab = “TDI”)*

11. Perform the flat-line test for a PhyloExpressionSet of step 3 and the TAI profile of step 7:

> *FlatLineTest (ExpressionSet = PhyloExpressionSetExample*,
>
> *permutations =10000*,
>
> *plotHistogram = TRUE)*

A. Perform the reductive hourglass test for a PhyloExpressionSet of step 3 and the TAI profile of step 7:

> *ReductiveHourglassTest (ExpressionSet = PhyloExpressionSetExample*,
>
> *modules = list(early = 1:2, mid = 3:5, late = 6:7)*,
>
> *permutations =10000*,
>
> *plotHistogram = TRUE)*

B. Perform the reductive early-conservation test for a PhyloExpressionSet of step 3 and the TAI profile of step 7:

> *EarlyConservationTest (ExpressionSet = PhyloExpressionSetExample*,
>
> *modules = list(early = 1:2, mid = 3:5, late = 6:7)*,
>
> *permutations =10000*,
>
> *plotHistogram = TRUE)*

12. Perform the flat-line test for a DivergenceExpressionSet of step 3 and the TDI profile of step 9:

> *FlatLineTest (ExpressionSet = DivergenceExpressionSetExample*,
>
> *permutations =10000*,
>
> *plotHistogram = TRUE)*

A. Perform the reductive hourglass test for a DivergenceExpressionSet of step 3 and the TDI profile of step 9:

> *ReductiveHourglassTest (ExpressionSet = DivergenceExpressionSetExample*,
>
> *modules = list(early = 1:2, mid = 3:5, late = 6:7)*,
>
> *permutations =10000*,
>
> *plotHistogram = TRUE)*

B. Perform the reductive early-conservation test for a DivergenceExpressionSet of step 3 and the TDI profile of step 9:

> *EarlyConservationTest (ExpressionSet = DivergenceExpressionSetExample*,
>
> *modules = list(early = 1:2, mid = 3:5, late = 6:7)*,
>
> *permutations =10000*,
>
> *plotHistogram = TRUE)*

13. Compute relative expression profiles for PS based on a PhyloExpressionSet of step 3:

> *REMatrix(PhyloExpressionSetExample)*

14. Visualize relative expression profiles for PS from the data of step 13:

> *PlotRE (ExpressionSet = PhyloExpressionSetExample*,
>
> *Groups = list(group = 1:12)*,
>
> *legendName = “PS”*,
>
> *xlab = “Ontogeny”*,
>
> *lty = 1*,
>
> *cex = 0.7*,
>
> *lwd = 5)*

A. Visualize relative expression profiles based on groups of PS from the data of step 13 (Fig. 4a):

> *PlotRE (ExpressionSet = PhyloExpressionSetExample*,
>
> *Groups = list(group_1 = 1:3, group_2 = 4:12)*,
>
> *legendName = “PS”*,
>
> *xlab = “Ontogeny”*,
>
> *lty = 1*,
>
> *cex = 0.7*,
>
> *lwd = 5)*

15. Compute relative expression profiles for DS based on a DivergenceExpressionSet of step 3:

> *REMatrix(DivergenceExpressionSetExample)*

16. Visualize relative expression profiles for DS based on the data from step 15:

> *PlotRE (ExpressionSet = DivergenceExpressionSetExample*,
>
> *Groups = list(group = 1:10)*,
>
> *legendName = “DS”*,
>
> *xlab = “Ontogeny”*,
>
> *lty = 1*,
>
> *cex = 0.7*,
>
> *lwd = 5)*

A. Visualize relative expression profiles based on groups of DS from the data of step 15 (Fig. 4b):

> *PlotRE (ExpressionSet = DivergenceExpressionSetExample*,
>
> *Groups = list(group_1 = 1:5, group_2 = 6:10)*,
>
> *legendName = “DS”*,
>
> *xlab = “Ontogeny”*,
>
> *lty = 1*,
>
> *cex = 0.7*,
>
> *lwd = 5)*

17. Quantify the statistical significance of group differences for PS based on the data of step 14A(Fig. 4c):

> *PlotBarRE (ExpressionSet = PhyloExpressionSetExample*,
>
> *Groups = list(group_1 = 1:3, group_2 = 4:12)*,
>
> *xlab = “Ontogeny”*,
>
> *ylab = “Mean Relative Expression”*,
>
> *cex =2)*

18. Quantify the statistical significance of group differences for DS based on the data of step 16A(Fig. 4c):

> *PlotBarRE (ExpressionSet = DivergenceExpressionSetExample*,
>
> *Groups = list(group_1 = 1:5, group_2 = 5:10)*,
>
> *xlab = “Ontogeny”*,
>
> *ylab = “Mean Relative Expression”*,
>
> *cex =2)*

### Protocol Timing

Computing the phylostratigraphic map can take several hours, several days, or even several weeks depending on the number of query sequences and the size of the database of proteins of completely sequenced genomes and may thus require a computing cluster, which can easily run these jobs in parallel, reducing the time to only a few hours.

Computing the divergence stratigraphic map takes 2 - 4 hours on an ordinary PC (step 2). All other steps (3 - 18) take less than 10 minutes on an ordinary PC. Analyses of other data sets might take more or less time depending on the number of samples, genes, and permutations. Computation time and command-input time for most steps in the protocol are only a few seconds. The most time-consuming steps are the computation of phylostratigraphic maps or divergence stratigraphic maps in cases where they are not yet available, data formatting for obtaining *PhyloExpressionSets* and *DivergenceExpressionSets*, or performing permutation tests if a large number of permutations is chosen.

### Protocol Results

Performing phylostratigraphy with Perl script <u>createPSmap.pl</u> and divergence stratigraphy with R package *orthologr* for the example data set (steps 1-2) yields a phylostratigraphic map (**Fig. 2**) and a divergence stratigraphic map for the studied species. Alternatively, users can input their own custom phylostratigraphic maps or divergence stratigraphic maps at this stage of the protocol (Supplementary Information). Matching these maps with gene expression data covering the studied developmental process (step 3) yields a PhyloExpressionSet (**Fig. 2a**) and a DivergenceExpressionSet (**Fig. 2b**). Applying steps 7-10 of the protocol to these two data sets yields TAI and TDI profiles for seven stages of *A.thaliana* embryo development (**Fig. 3**).

The *flat-line test* (steps 11-12) yields *p*-values of 1.2e-09 (TAI) and 8.8e-06 (TDI), stating that both profiles deviate significantly from a flat line. The *reductive hourglass test* (steps 11.A and 12.A) yields *p*-values of 8.0e-09 (TAI) and 4.8e-04 (TDI), stating that both profiles are compatible with an hourglass pattern. The *reductive early-conservation test* (steps 11.B and 12.B) yields *p*-values of 0.99 (TAI) and 0.96 (TDI), stating that both profiles deviate significantly from an early-conservation pattern.

Steps 13-16 yield relative expression profiles for all PS (**Fig. 4a**) and DS (**Fig. 4b**). For the example data set, we observe that evolutionarily conserved genes (genes of PS 1-3 that emerged before embryogenesis) and evolutionarily young genes (genes of PS 4-12 that emerged after embryogenesis) show qualitatively different relative expression profiles. Figure 4a shows that high PS have similar relative expression profiles, whereas low PS have relative expression profiles that are dissimilar to each other and dissimilar to those of high PS. Figure 4b shows that DS 2-10 have similar relative expression profiles, whereas DS 1 shows an antagonistic relative expression profile.

Applying the Kruskal-Wallis rank-sum test to the observed relative expression levels of the two groups of PS (step 17) and the two groups of DS (step 18) yields *p*-values < 0.05 in both cases, stating that the observed differences of the relative expression levels between low and high PS and between low and high DS are statistically significant (**Fig. 4c**).

Together, this protocol allows users to perform phylotranscriptomic analyses of biological processes of their interest in a standardized manner. The intuitive adaptability of the protocol to any species and any biological process is achieved by the modular structure of the protocol, Perl script createPSmap.pl, and R packages *orthologr* and *myTAI*, which provide an open-source implementation of the protocol. The Perl script and both R packages can be used for performing phylotranscriptomic analyses in a reproducible manner. The documentation of each function as well as the tutorials included in the Supplementary Information provide additional details of the functionality provided by createPSmap.pl, *orthologr*, and *myTAI* so that this protocol can be easily applied by a broad set of users to numerous of phylotranscriptomic studies of various biological processes in the future.

## Author Contributions

HGD conceived the protocol. HGD, AG, TDL, MQ, and IG designed the protocol. HGD designed and implemented *orthologr* and *myTAI*. AG designed and implemented *<u>createPSmap.pl</u>*. HGD, TDL, MQ, and IG wrote the manuscript.

## Acknowledgments

We thank Jan Grau, Anne Hoffmann, Philipp Janitza, Kristian Karsten Ullrich, Sarah Scharfenberg Claus Weinholdt, and Sebastian Wussow for valuable discussions and Deutsche Forschungsgemeinschaft (grants no. FZT 118, GR 3526/2, GR 3526/6, GR 3526/7, GR 3526/8, Qu 141/5, Qu 141/6, and Qu 141/7) and SKW Piesteritz for financial support.

